# CASC: Content-Aware Compaction of Sparse Microscopy Images

**DOI:** 10.64898/2026.06.15.731850

**Authors:** Gail McConnell

## Abstract

Microscopy datasets are often spatially sparse, wherein relevant structures occupy only a small fraction of the total field of view (FOV), leaving large regions of background devoid of signal. This inherent inefficiency creates file sizes that are larger than needed, which increases the time needed for computational image data analysis and processing, and means unnecessarily large data volumes. In this work, a classical open-source method for content-aware spatial compaction of microscopy images (CASC) is reported that explicitly removes spatial redundancy by reorganising foreground objects into a new, smaller image. CASC combines adaptive intensity normalisation, statistical thresholding, morphological refinement, and connected-component analysis to isolate foreground structures. These structures are then extracted with contextual padding and repacked into a compact domain using a heuristic shelf-based spatial packing strategy. CASC intentionally destroys the spatial topology of the image but preserves pixel intensities exactly, retaining object-level information. The method achieves demonstrable reductions in image area and background content while maintaining high object-level preservation of biological structures, with a reduction in file size of more than 390-fold shown in real image datasets.

## 1. Introduction

High-resolution microscopy has become ubiquitous in modern biological and biomedical research, enabling the visualisation and measurement of cellular and sub-cellular structural detail with increasing precision. However, many microscopy datasets remain highly spatially sparse, with biologically relevant structures occupying only a small fraction of the field of view (FOV) (1). In such datasets, cells, nuclei, or labelled organelles may be distributed across large image areas while most pixels correspond to background or unstained specimen regions. As image dimensions increase, operations such as thresholding, segmentation, visualisation, storage, and data transfer must still process the full image domain despite the limited spatial occupancy of meaningful signal (2). This results in inflated data volumes and inefficient use of computational resources throughout image acquisition, analysis, storage, and sharing. The problem is particularly pronounced in mesoscale microscopy, where multi-millimetre or centimetre-scale FOVs may contain sparse foreground structures distributed across extremely large image areas (3–6).

Traditional approaches to reducing microscopy data size rely on image compression algorithms that exploit redundancy in pixel values while preserving the spatial topology of the image (7,8). Although such methods reduce storage requirements, they do not address the spatial inefficiency of the image representation itself. Manual cropping of regions of interest offers a partial solution, but sparse objects may be distributed throughout the image, making exhaustive cropping impractical for large datasets and potentially introducing user bias into downstream analysis.

An alternative approach is presented here. Rather than treating the image as a fixed spatial grid of pixels, the image can instead be interpreted as a collection of discrete foreground objects embedded within space. Under this interpretation, empty image regions may be removed entirely and foreground objects reorganised into a more compact representation. Here, compaction refers to the spatial rearrangement of segmented foreground objects into a reduced image domain while preserving their original pixel values. Unlike conventional image compression methods, which preserve spatial topology while reducing pixel redundancy, this approach explicitly treats spatial arrangement itself as a removable source of redundancy. While object pixel intensities can be preserved exactly, rearrangement of objects inevitably alters local and global spatial relationships. It is therefore essential not only to perform compaction efficiently, but also to quantify the extent to which spatial structure is preserved or destroyed.

In this work, a complete pipeline for content-aware compaction of sparse microscopy images (CASC) is presented for object-centric datasets in which biologically relevant information is primarily associated with individual foreground structures rather than their absolute spatial arrangement. CASC combines foreground segmentation, connected-component extraction, and heuristic spatial packing to reduce dead space and minimise spatial redundancy while preserving object-local morphology and intensity information. Conceptually, CASC reframes sparse microscopy images not as fixed spatial entities, but as collections of independent foreground objects that can be reorganised into a more compact and computationally efficient dataset for processing, sharing, storing, and archiving.

## 2. Methods

The CASC pipeline was implemented in Python and executed on a laptop computer running Microsoft Windows 11 Enterprise (version 10.0.26200 Build 26200) on an x64-based architecture. The system was equipped with an Intel® Core™ Ultra 5 135U processor (12 cores, 14 logical processors) and 16 GB RAM. GPU acceleration was not used during image processing or analysis.

The software implementation used open-source Python libraries including NumPy (9), SciPy (10), scikit-image (11), pandas, Pillow, and tifffile for image processing, segmentation, connected-component analysis, metric computation, and file input/output operations. The source code for the method is available at the GitHub resource: https://github.com/gailmcconnell/CASC.

### 2.1 Image Representation and Preprocessing

The proposed methodology of CASC is shown schematically in Figure 1. The approach consists of a sequence of transformations that progressively convert an input microscopy image into a compact representation while preserving object-level information. The pipeline begins with image normalisation and foreground detection, followed by connected-component extraction, spatial packing, and reconstruction of the compact image. Finally, a set of evaluation metrics is computed to assess the object-level preservation of the transformation.

**Figure 1.**
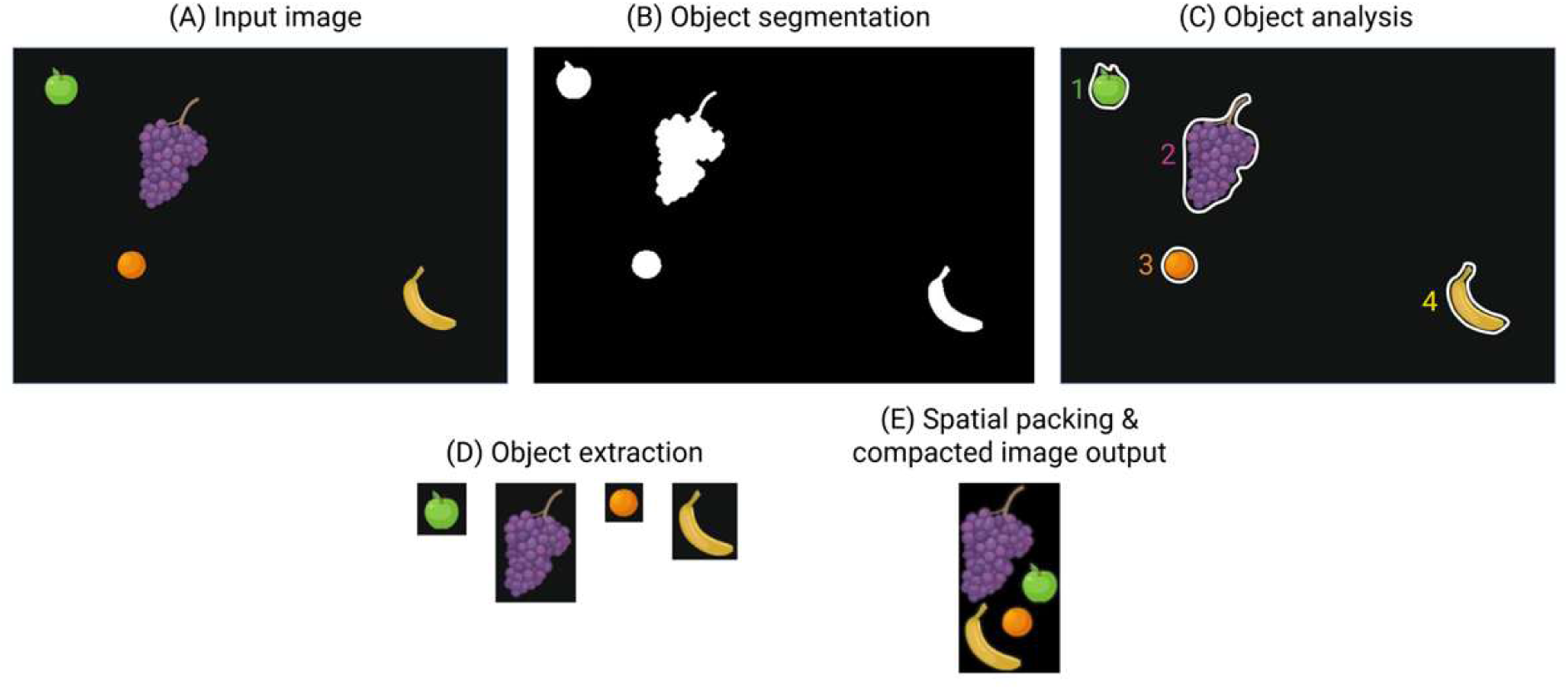
Schematic overview of the content-aware spatial compaction (CASC) pipeline. (A) Input image containing sparsely distributed objects within a large field of view. (B) Thresholding and segmentation are used to distinguish objects from the background, generating a binary mask. (C) Connected-component analysis identifies and labels individual foreground objects. (D) Each detected object is extracted as an individual image crop with associated spatial information recorded. (E) Extracted objects are spatially repacked into a compact non-overlapping arrangement to generate a reduced-area output image while preserving the original object pixel intensities. Figure 1(A) was created using BioRender.

To ensure generality, the method first converts the input image into a two-dimensional (2D) representation suitable for foreground detection. For greyscale images, this representation is used directly. For colour images, a luminance transformation is applied, producing a weighted combination of the red, green, and blue channels. The resulting detection image was converted to floating-point representation and normalised to the range [0,1].

### 2.2 Foreground Detection

Foreground detection was performed using a multi-stage preprocessing and thresholding workflow. Local contrast was first enhanced using contrast-limited adaptive histogram equalisation (12), and foreground regions were then identified using global Otsu thresholding (13). To accommodate images in which foreground objects appeared darker than the background, the binary mask was automatically inverted when the initial foreground fraction exceeded 50% of the image area.

Morphological refinement was subsequently applied to improve mask quality and suppress segmentation artefacts. Binary opening and closing operations were performed using disk-shaped structuring elements with radii of 1 and 2 pixels, respectively. Connected regions below a minimum object area threshold of 25 pixels were removed, and small internal holes were filled using an area threshold of either 64 pixels or twice the minimum object area, whichever was larger.

The resulting binary mask was used to define candidate foreground regions for downstream connected-component extraction and spatial packing. Separate masks were additionally generated for collision-aware object placement and visual rendering operations during image reconstruction.

### 2.3 Connected Component Extraction

Once a clean binary mask has been obtained, connected-component analysis is used to identify individual foreground objects (14). Each connected component is defined as a maximal set of foreground pixels that are connected under an eight-neighbourhood adjacency criterion. For each component, geometric properties such as area, bounding box, and centroid are computed.

To preserve contextual information, the bounding box of each component is expanded by a fixed padding margin (default = 8 pixels). This padding ensures that the extracted region includes not only the object itself but also a small surrounding area, which may be important for interpretation or subsequent analysis. The padded region is then used to extract both the corresponding image patch and a binary mask indicating the pixels belonging to the component. Each component is represented as an object containing its original spatial coordinates, geometric properties, and pixel data.

### 2.4 Spatial Packing

The problem of arranging the extracted components into a compact layout can be viewed as fitting a collection of rectangular regions into a smaller image area without allowing them to overlap. This corresponds to a two-dimensional rectangle packing problem, which is computationally challenging because the number of possible arrangements increases rapidly with object number (15–17). As a result, exhaustively testing all possible layouts is not practical for typical datasets. Instead, a fast and reliable approximation strategy based on a shelf-style packing approach is used (16). This produces an efficient and well-organised layout without requiring excessive computation.

In the shelf packing scheme, components are first sorted in descending order of height (i.e. pixels along the y-axis of the original image). They are then placed sequentially along horizontal rows, or “shelves,” within the output canvas. When the addition of a component would exceed the height of the current row, a new row is started below the previous one. The height of each row is determined by the tallest component it contains, and the overall height of the canvas is the sum of the row heights.

The width of the canvas is estimated based on the total area of the components. While the shelf algorithm does not guarantee optimal packing, it offers a favourable balance between computational efficiency and packing quality, making it well suited to large image data.

### 2.5 Image Reconstruction

Once the placement of each component has been determined, a new image canvas is initialised with dimensions corresponding to the packed layout. The canvas is filled with zeros, representing background. Each component is then placed onto the canvas at its assigned location, using its binary mask to ensure that only foreground pixels are copied. For multi-channel images, this operation is performed independently for each channel.

Importantly, no interpolation or transformation of pixel values is performed during this process. The pixel intensities of each component are copied exactly from the original image, ensuring that the compact representation preserves the intensity of objects. The only change introduced by the method is the spatial rearrangement of components.

### 2.6 Evaluation Metrics

To quantify the effects of spatial compaction, a set of object-level metrics are recorded. The reduction in FOV area is measured as the relative decrease in total pixel count between the original and output images. Foreground retention is defined as the fraction of foreground pixels preserved after compaction, which should ideally be equal to one given that pixel values are copied exactly.

To assess changes in spatial relationships, the centroids of the components before and after packing are recorded. The preservation of local structure is evaluated by comparing the sets of nearest neighbours for each component using a Jaccard similarity measure (18). Global structure is assessed using the Spearman rank correlation between pairwise distance matrices, which captures the extent to which relative distances are preserved. Finally, local geometric distortion is quantified as the relative change in nearest-neighbour distances. These metrics provide a comprehensive view of the trade-offs inherent in spatial compaction, allowing the method to be evaluated in a principled manner.

### 2.7. Specimens and specimen preparation

*Escherichia coli* strain JM105-miniTn7-*gfp* was grown in lysogeny broth at 37 °C with shaking at 200 rpm to mid-exponential phase. Cultures were diluted to 1:10000 for imaging using widefield epifluorescence microscopy as described below. For microscopy, a thin monolayer of cells was prepared on Type 1.5 glass coverslips. A 10 µL drop of the planktonic bacterial culture was applied to the coverslip and allowed to settle for several minutes to facilitate surface attachment. Excess liquid was gently removed, and the prepared sample was mounted onto a microscope slide and sealed to prevent evaporation.

HeLa cells were cultured in Dulbecco’s modified Eagle medium plus GlutaMAX medium (Gibco) supplemented with 10% (v/v) FBS and 100 μg/ml penicillin-streptomycin. Cells were plated onto Type 1.5 coverslips a day prior to fixation, then fixed with 4% paraformaldehyde for 15 min at 37°C, followed by a permeabilisation step with 2.5% FBS and 0.3% Triton X-100 in PBS for 30 min at room temperature (19). A paxillin monoclonal antibody (5H11, 1:500) was conjugated to a fluorescent secondary antibody (Alexa Fluor Plus 594, Life Technologies) was applied to the fixed cell specimen as per the manufacturer instructions. Specimens were mounted in ProLong Glass Antifade mountant (Life Technologies) and mounted onto microscope slides for imaging.

A proprietary specimen of bovine pulmonary artery endothelial (BPAE) cells (FluoCells™ Prepared Slide #1, Thermo Fisher Scientific) labelled with MitoTracker Red CMXRos, Alexa Fluor 488 Phalloidin, and DAPI was also imaged.

### 2.8. Imaging

Fluorescence microscopic imaging of the *E. coli* cell specimens was performed using a widefield epifluorescence microscope (Ti2 microscope, Nikon) equipped with a monochrome Prime BSI scientific CMOS camera (Photometrics) controlled by proprietary software (NIS Elements, Nikon). Images were acquired using a 20x/0.7 numerical aperture objective lens. A light emitting diode engine (pE-4000, CoolLED) was used for excitation of fluorescence with a 490 nm source. Images were obtained at a camera exposure time of 100 ms, with 11-bit images recorded. Image data were recorded in the OME TIFF format.

Fluorescence mesoscale imaging of the BPAE cell specimens was performed using a Mesolens (Mesolens Ltd) equipped with a monochrome large format camera (MX2457MR-SY-X4G4G3-FF, Ximea) controlled by proprietary software (CamTool, Ximea). Images were acquired using a light emitting diode engine (pE-4000, CoolLED) for excitation of fluorescence with a 385 nm source. Images were obtained at a camera exposure time of 50 ms, with no additional gain and 8-bit images were recorded in the OME TIFF format.

Fluorescence mesoscale imaging of the HeLa cell specimens was performed using a Mesolens (Mesolens Ltd) equipped with a chip-shifting camera (VNP-29MC, Vieworks) controlled by open-source software (MesoCam). With a 3×3 chip-shift, the sampling rate was 4.46 px/µm, corresponding to a 224 nm pixel size, satisfying Nyquist sampling. Images were acquired using a light emitting diode engine (pE-4000, CoolLED) for excitation of fluorescence with a 594 nm source. Images were obtained at a camera exposure time of 200 ms, with no additional gain and 8-bit images were recorded in the OME TIFF format.

### 2.9. Application of CASC to image data

Images of the *E. coli*, BPAE, and HeLa cell specimens were used with the CASC Python code to produce the new compacted image outputs. For presentation of data, the colour look-up table of the images were changed, with the nuclei shown with the NanoJ-Orange colour look up table and the *E. coli* cells shown with the Fire look up table. Both colour look-up tables were applied in FIJI (20).

### 2.10 Simulation of noise and contrast conditions

To assess robustness of CASC with image quality variation, synthetic noise and contrast perturbations were applied using custom Python scripts.

Image degradation by noise was simulated using additive Gaussian noise implemented in Python (OpenCV, NumPy). The input image was converted to 8-bit format, and a sequence of N=16 images was generated with progressively increasing noise levels. Noise magnitude was controlled by varying the standard deviation (σ) of zero-mean Gaussian noise according to σ=σ _max_·(i/(N-1))², where σ_max_ = 140. This quadratic scaling resulted in finer sampling at low noise levels and coarser sampling at high noise levels. Noise was applied uniformly across the image. The noisy image was computed as a weighted combination of the original image and the generated noise.

For each noise level, object detection was performed using thresholding followed by watershed segmentation. Objects smaller than 20 pixels were excluded. The number of detected objects was recorded as a function of σ.

To assess the robustness of CASC to image contrast, the image of DAPI-stained BPAE cell nuclei was cropped in size to produce a smaller region of interest of 360 µm x 360 µm. The contrast of this image was systematically reduced using a linear intensity-compression procedure implemented in Python. Minimum and maximum pixel intensities were computed (per channel for colour images), and a sequence of 16 images was generated by progressively compressing the intensity range. For each image i, a contrast compression factor α was defined as α=i/(N-1), where N=16 is the number of images.

The target intensity range was linearly interpolated between the original range (α=0) and a compressed range of 0 to 1 counts (α=1). Pixel intensities were remapped accordingly using a linear transformation, clipped to the range 0-255, and these were saved as 8-bit TIFF images.

For each contrast level, object detection was performed using thresholding followed by watershed segmentation. Small objects (<20 pixels) were excluded. The number of segmented objects was recorded for each image to quantify the effect of contrast reduction on detection performance.

All noise- and contrast-modified images were then processed using the same compaction pipeline. Outputs and evaluation metrics were generated for each case, allowing method performance to be assessed under progressively degraded image conditions.

### 2.11 Analysis of scalability

Computational scaling behaviour of CASC was evaluated using synthetic benchmark datasets generated with custom Python scripts. The effects of image dimensions and object density on processing time were measured.

To evaluate scaling with image size independently of object number, synthetic TIFF images containing a single circular object were generated at image sizes ranging from 32 pixel x 32 pixel to 16384 pixel x 16384 pixel.

To evaluate scaling with object density, a further synthetic dataset was generated. This dataset contained increasing numbers of non-overlapping circular objects placed randomly throughout the image field. An image size of 16384 pixel x 16384 pixel was chosen to mimic mesoscale imaging datasets, with the number of objects increasing from 1 to 2048, thus emulating sparse datasets.

In both cases, processing time measurements represented total end-to-end runtime for the complete CASC workflow from pipeline initiation to completion. Runtime was measured using wall-clock timing implemented with Python time.time().

### 2.12 Comparison with image compression methods

The compacted outputs of CASC applied to the image datasets described in 2.9 were benchmarked against conventional image compression approaches. Raw image datasets were converted to lossy JPEG and lossless PNG, ZIP, JPEG 2000, and ZARR (8) formats using standard encoding settings. For each method, the resulting file sizes were recorded.

### 2.13 Pixel-preserving montage generation

To provide a comparison with a conventional object-based visualisation approach, pixel-preserving montages were generated from the same connected components used as inputs to CASC. Following connected-component extraction, each segmented object was cropped using its padded bounding box as described in Section 2.3. No interpolation, scaling, rotation, or intensity transformation was applied. Consequently, each crop retained the original pixel dimensions and intensity values of the corresponding object.

The extracted object crops were arranged using a shelf-based montage layout. Crops were first sorted in descending order of height and then placed sequentially along horizontal rows. When the addition of a crop would exceed the target montage width, a new row was initiated. A fixed padding margin was maintained between adjacent crops to avoid overlap. Unlike CASC, which reconstructs the output image using object masks, the montage representation retained the complete rectangular bounding box associated with each extracted crop, including any background pixels contained within the bounding region.

The dimensions of the montage canvas were determined from the total area of the extracted crops and the resulting shelf arrangement. The final montage therefore preserved object morphology, pixel dimensions, and intensity values exactly, while discarding the original spatial arrangement of the objects. Because the montage retained rectangular crop boundaries and inter-object spacing, it provided a useful benchmark for assessing the extent to which additional reductions in spatial redundancy could be achieved through the object-aware packing and reconstruction strategy employed by CASC. This comparison was included to determine whether the reduction in spatial redundancy achieved by CASC could be replicated using a simpler object-rearrangement strategy that preserves object dimensions and pixel intensities.

## 3. Results

### 3.1 Compaction of BPAE nuclei images with CASC

Figure 2 shows the result of CASC applied to the widefield epifluorescence mesoscopy image of DAPI-stained nuclei in a BPAE cell specimen. The original image, shown in Figure 2(A) had a file size of 37.4 MB file size for the full image. Applying CASC to the image shown in Figure 2(A) produced the compacted output shown in Figure 2(B). This image retained object level information without loss of pixel intensity for each object, and had a filesize of 731 kB, corresponding to a 51-fold reduction in data volume compared to the original image.

**Figure 2.**
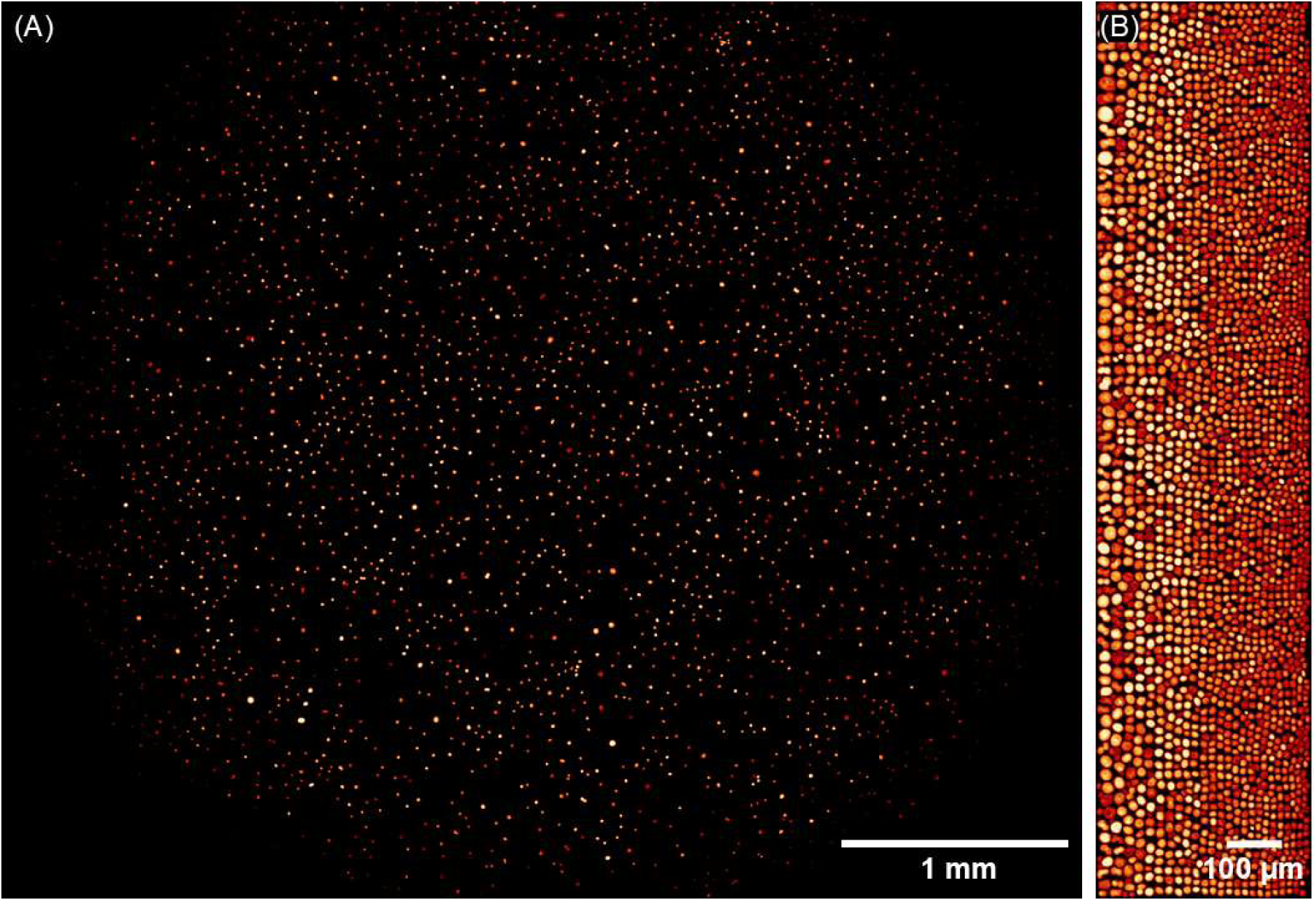
Representative images of DAPI-stained BPAE cell nuclei before and after compaction with CASC. (A) Widefield epifluorescence mesoscopy image of DAPI-stained BPAE cell nuclei obtained with the Mesolens, showing sparse and heterogeneous organisation across the field of view. (B) Nuclei following compaction via CASC, preserving object morphology and intensity values. A reduction in file size from 37.4 MB for the full image presented in (A) to 731 kB for the output shown in (B), corresponding to a 51-fold reduction in data volume. Using distance transform-based watershed segmentation to separate touching features and counts objects, the compacted output retains 96.5% of the objects identified in the original image. Scale bars: (A) 1 mm, (B) 100 µm.

Using distance transform-based watershed segmentation in FIJI to separate touching features and count objects, a total of 2796 nuclei were counted in the original image compared to 2699 nuclei in the compacted output, corresponding to a 96.5% preservation of the objects identified in the original image.

Quantitative analysis of this dataset demonstrated extensive reorganisation of object spatial relationships following content-aware compaction. Nearest-neighbour retention measured using Jaccard similarity was effectively absent (Jaccard similarity = 0.0004), indicating that almost none of the original local neighbourhood relationships were preserved after repacking. Similarly, the pairwise distance rank correlation was close to zero and slightly positive (Spearman ρ = 0.004), demonstrating that the global ordering of pairwise object distances was not retained in the compacted representation. The mean relative nearest-neighbour distortion was 11.374, indicating large alterations in local geometric spacing between neighbouring objects following rearrangement within the compacted image.

### 3.2 Compaction of *E. coli* images with CASC

Figure 3 shows a representative image of *E. coli* JM105-miniTn7-gfp cells before and after compaction with CASC. Figure 3(A) shows a raw widefield epifluorescence image of *E. coli* JM105-miniTn7-gfp cells showing a sparse cell population across the image. Yellow, green, and cyan boxes highlight different regions of interest (ROIs) containing cells. A ‘Fire’ look-up table has been applied to the raw greyscale image. Figure 3(B) displays the result of CASC applied to the image shown in Figure 3(A). A reduction in file size from 8.2 MB for the full image presented in Figure 3(A) to 21.0 kB for the output shown in Figure 3(B) was measured, corresponding to a 390-fold reduction in data volume.

**Figure 3.**
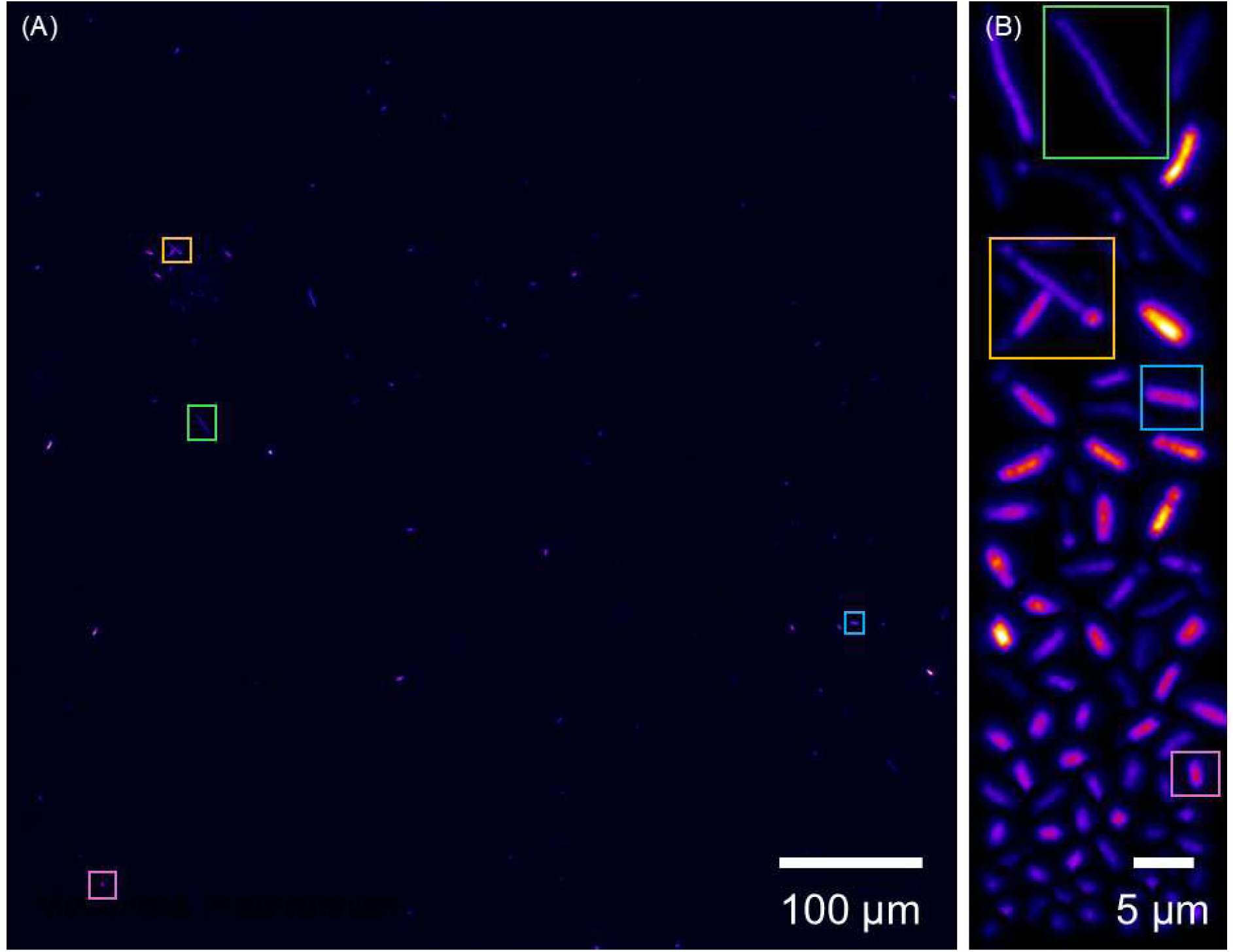
Representative images of *E. coli* JM105-miniTn7-*gfp* cells before and after compaction with CASC. (A) Widefield epifluorescence image of *E. coli* JM105-miniTn7-*gfp* cells showing sparse and heterogeneous organisation across the field of view. Yellow, green, and cyan boxes highlight different regions of interest (ROIs) containing cells. A ‘Fire’ look-up table has been applied to the raw greyscale image. (B) Compaction of (A) performed with CASC, showing *E. coli* cells with object morphology and intensity values preserved. A reduction in file size from 8.20 MB for the full image presented in (A) to 21.0 kB for the output shown in (B), corresponding to a 390-fold reduction in data volume. Scale bars: (A) 100 µm, (B) 5 µm.

Quantitative analysis of this dataset demonstrated extensive reorganisation of object spatial relationships following content-aware compaction. Nearest-neighbour retention measured using Jaccard similarity remained absent (Jaccard similarity = 0.00), indicating that none of the original local neighbourhood relationships were preserved after repacking. Similarly, the pairwise distance rank correlation was close to zero and slightly negative (Spearman ρ = - 0.02), demonstrating that the global ordering of pairwise object distances was not retained in the compacted output. The mean relative nearest-neighbour distortion was 0.83, indicating some alteration of local geometric spacing between neighbouring objects following rearrangement within the compacted image.

### 3.3 Preservation of object morphology distributions with CASC

To evaluate whether CASC preserved object-level morphological statistics, segmented object areas were compared between the original and compacted images using a fluorescence mesoscopy image dataset of paxillin-labelled HeLa cells. Data are shown in Figure 4.

**Figure 4.**
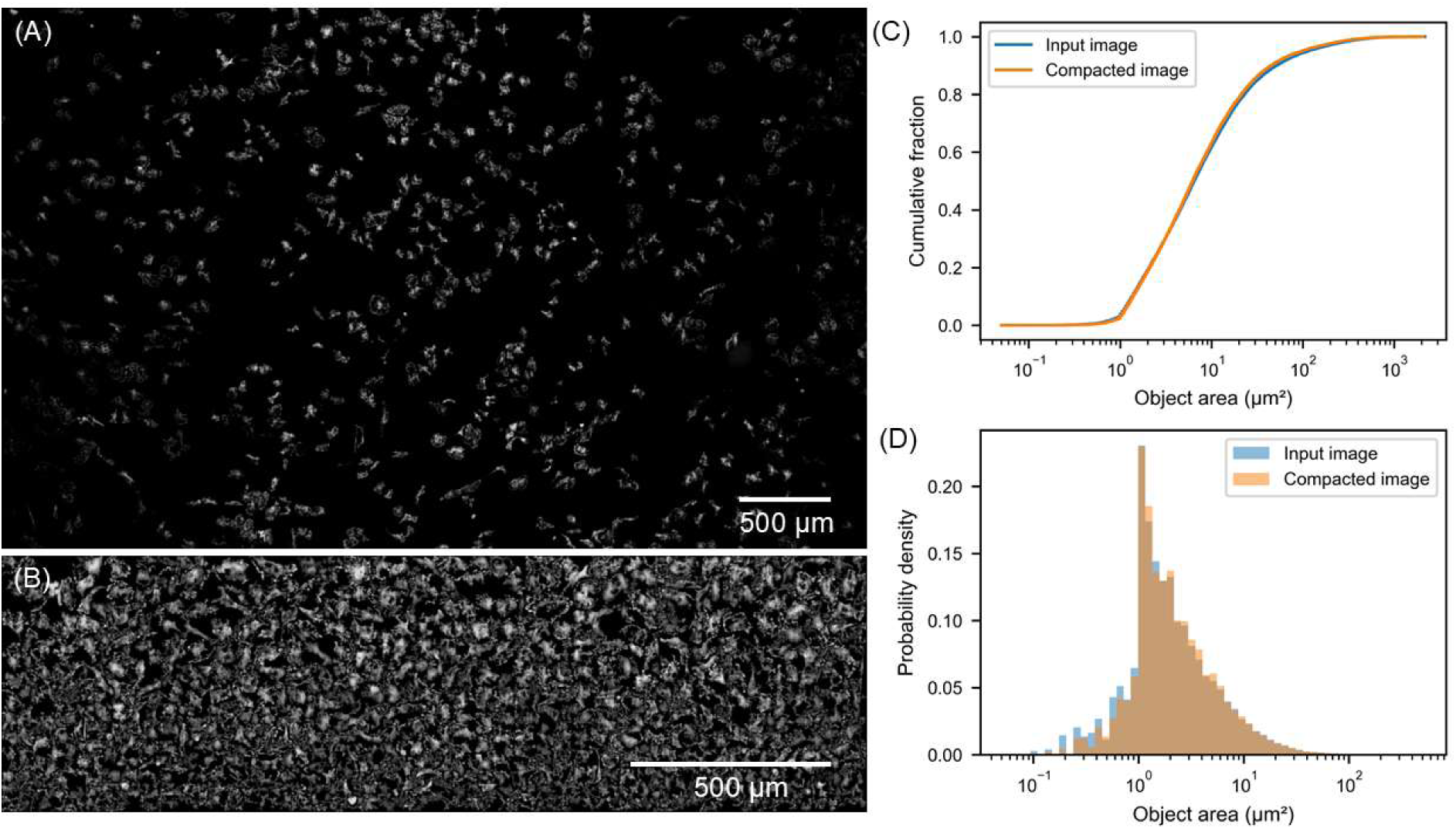
Preservation of object morphology distributions following content-aware compaction with CASC. (A) Raw input fluorescence mesoscopy image of HeLa cells labelled with a fluorescent paxillin stain, showing sparse spatial distribution of objects across the field of view. (B) Corresponding compacted image generated using the CASC pipeline. (C) Empirical cumulative distribution functions (ECDFs) of segmented object areas measured in the input and compacted images. Strong overlap between the distributions demonstrates preservation of cumulative object morphology statistics following compaction. (D) Log-scaled probability density histograms of object areas for the input and compacted images. Quantitative analysis identified 23022 objects in the input image and 22509 objects in the compacted image, corresponding to 97.8% object retention.

The compacted image, shown in Figure 4(B), retained the same foreground signal content as the input image, with the apparent increase in density resulting solely from repacking of objects into a smaller spatial domain. Quantitative analysis identified 23022 objects in the input image and 22509 objects in the compacted image, corresponding to 97.8% object retention following segmentation and compaction. The median object area changed from 6.47 µm² in the original image to 6.22 µm² in the compacted image, representing a median shift of -3.88%.

Empirical cumulative distribution functions (ECDFs) and log-scaled probability density histograms demonstrated strong overlap between the object area distributions measured before and after compaction. Statistical comparison using the Kolmogorov-Smirnov test produced a KS statistic of 0.0182, indicating only minor differences between the distributions despite the large number of measured objects.

These results demonstrate that CASC preserves global object morphology statistics with high fidelity while substantially reducing spatial redundancy within the image representation.

### 3.4 Effect of image noise on CASC

Figure 5 shows the effect of progressively increasing Gaussian noise on object detection performance to data used with CASC. At low and moderate noise levels, the number of detected objects remained relatively stable, indicating that the foreground detection and segmentation stages were robust to moderate image degradation. This suggests that the preprocessing and thresholding operations retained sufficient foreground-background separability despite increasing pixel-level noise.

**Figure 5.**
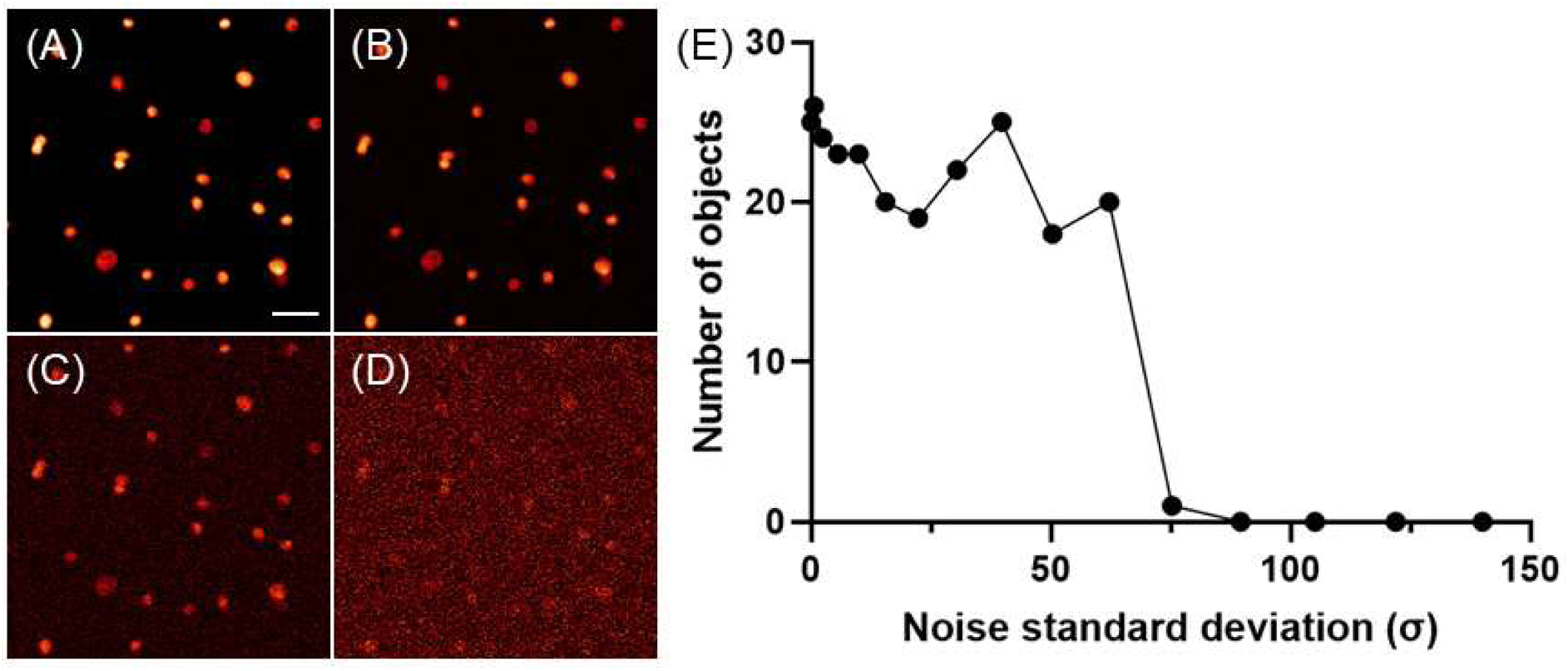
The effect of noise on content-aware compaction with CASC. Input images with increasing Gaussian noise standard deviation (A) σ = 0, (B) σ = 5.60, (C) σ = 30.49, and (D) σ = 75.3. (E) Number of detected objects using CASC as a function of Gaussian noise standard deviation following thresholding and watershed segmentation. Object detection remained relatively stable at low-to-moderate noise levels before collapsing rapidly at high noise levels. Scale bar: 50 µm.

As noise levels increased further, object counts declined progressively, followed by rapid segmentation failure at the highest noise conditions. This behaviour is consistent with deterioration of object boundary definition and increasing fragmentation of foreground regions, which impaired subsequent watershed-based separation of objects.

Importantly, the observed degradation originated primarily from failures in foreground detection and segmentation rather than from the spatial packing stage itself. Once objects were successfully segmented, compaction and reconstruction preserved object morphology and pixel intensities without additional degradation. These results therefore indicate that robustness of the overall CASC pipeline is governed principally by the quality of the initial segmentation stage.

### 3.5 Effect of image contrast on CASC

Figure 6 shows the effect of progressive contrast compression on object detection performance when using CASC. At low and moderate levels of contrast reduction, object counts remained largely stable, demonstrating tolerance of the segmentation workflow to moderate reductions in dynamic range.

**Figure 6.**
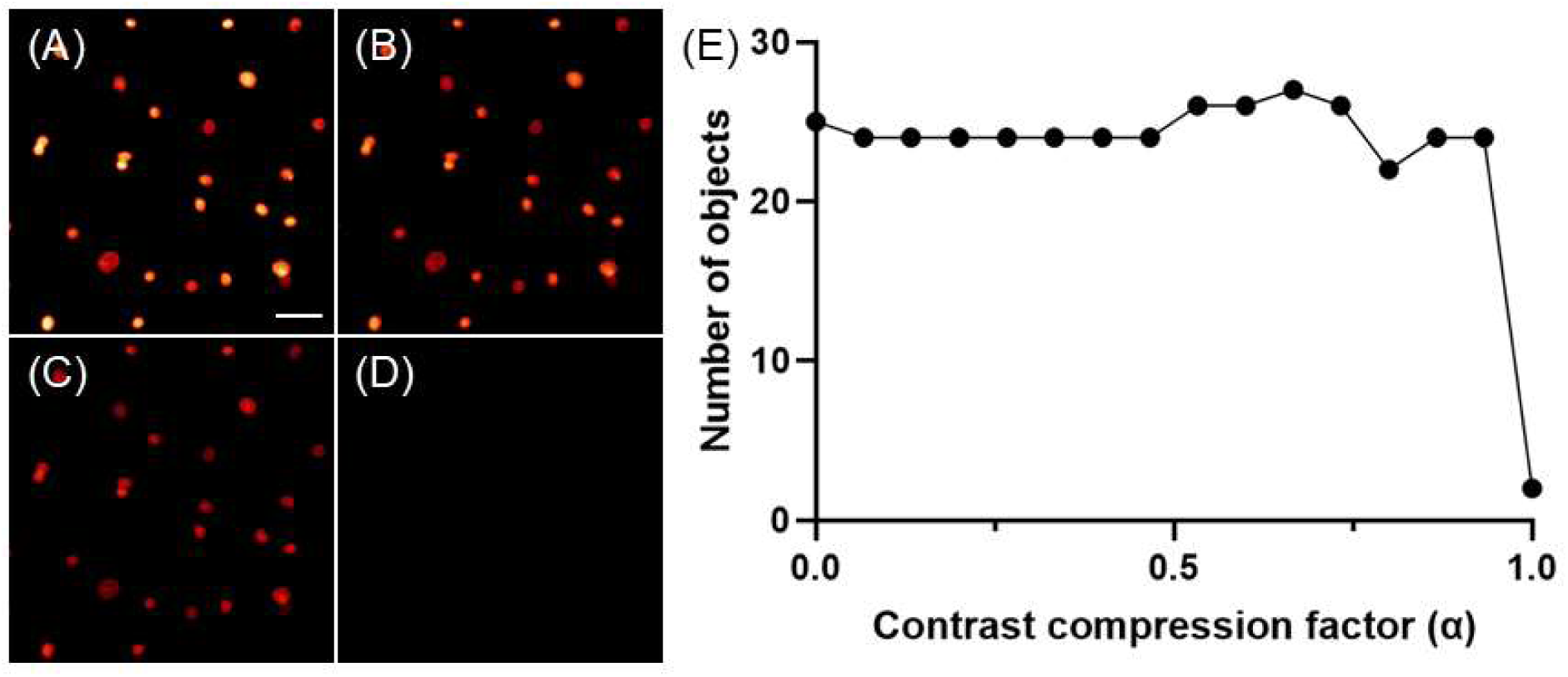
The effect of image contrast on content-aware compaction with CASC. Representative input images following progressive contrast compression are shown for contrast compression factors (A) α = 0, (B) α = 0.33, (C) α = 0.67, and (D) α = 1.0. (E) Number of detected objects as a function of contrast compression factor following thresholding and watershed segmentation. Object detection remained relatively stable across moderate contrast compression levels before collapsing at very low contrast (α = 1.0). Scale bar: 50 µm.

However, object detection performance deteriorated rapidly at high levels of contrast compression. Under these conditions, foreground and background intensities became increasingly indistinguishable, reducing the effectiveness of global thresholding and leading to merging and disappearance of segmented objects.

Compared with additive noise perturbation, contrast compression produced a more abrupt collapse in detection performance, indicating that the segmentation pipeline was more sensitive to loss of global intensity separation than to stochastic pixel-level fluctuations. This behaviour is consistent with the dependence of Otsu thresholding on separability between foreground and background intensity distributions.

### 3.6 Scalability of CASC

Computational benchmarking demonstrated distinct scaling behaviours associated with image dimensions and object density. These results are shown in Figure 7.

**Figure 7.**
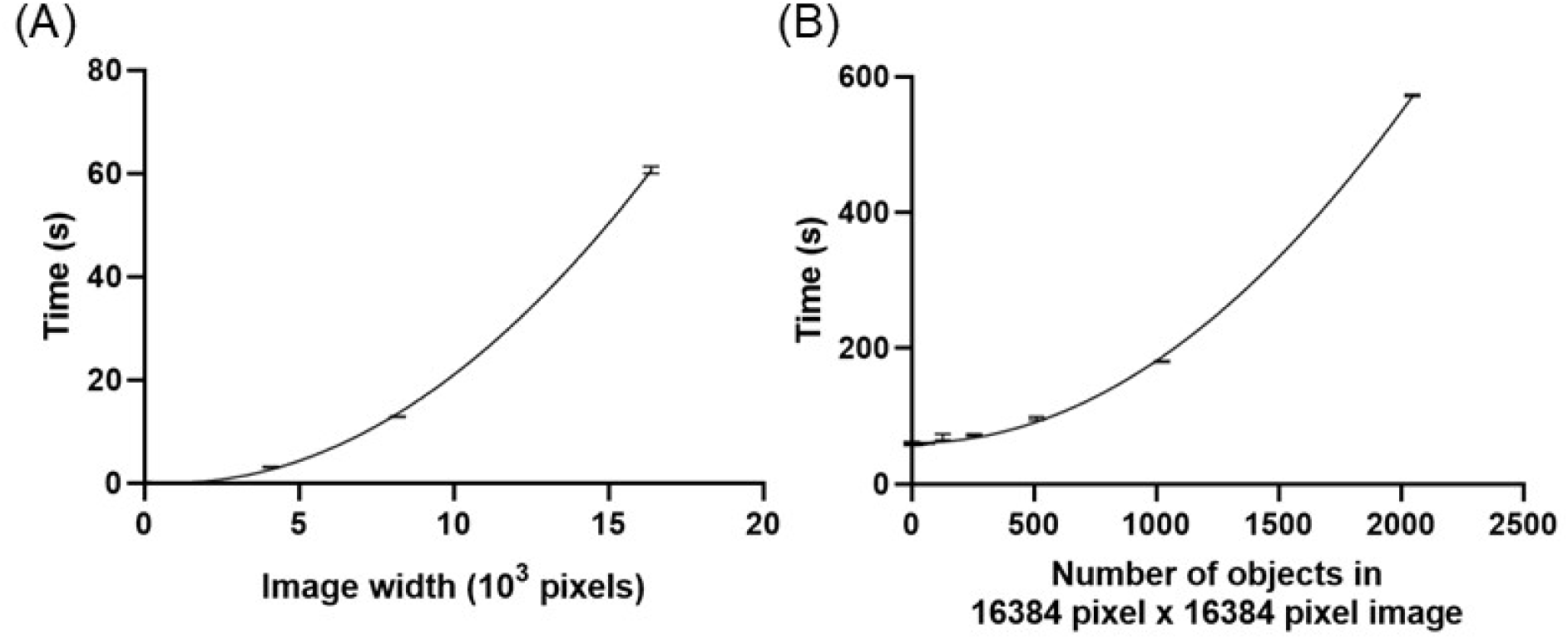
The effect of image size and number of objects on processing time for compaction with CASC. (A) Mean end-to-end processing time as a function of image size for synthetic images containing a single foreground object. The runtime scaled approximately with total image pixel count. (B) Mean processing time as a function of object number in 16384 pixel x 16384 pixel images. The runtime scaled nonlinearly with the object number. Error bars represent the standard deviation across n=5 measurements, and trendlines indicate empirical regression fits.

When object number was held constant at a single foreground object per image, the runtime of CASC increased nonlinearly with image dimensions, as shown in Figure 7(A). Runtime remained below 1 s for smaller images but increased rapidly at larger image sizes, reaching approximately 13 s for 8192 pixel x 8192 pixel images, and approximately 60 s for 16384 pixel x 16384 pixel images. The runtime scaled approximately with total pixel number, which is consistent with computational cost being dominated by operations that scale with total pixel count, including thresholding and occupancy-mask generation.

At fixed image dimensions of 16384 pixel x 16384 pixel but varying the number of objects in the image, the runtime of CASC increased nonlinearly, as shown in Figure 7(B). Runtime increased from a few seconds for sparse images to approximately 575 s for images containing 2048 objects. The empirical trend was consistent with nonlinear growth in processing time at high densities, indicating that object-packing operations become increasingly expensive as occupancy within the image increases.

Collectively, these results demonstrate that the computational cost of CASC depends strongly on both image dimensions and object density, with different stages of the compaction pipeline dominating under different conditions. For sparse datasets, runtime is governed primarily by image-wide preprocessing operations that scale with total pixel count. At high object densities, however, irregular object-packing operations become the main bottleneck. As such, CASC is best applied to sparse image datasets.

### 3.7 Comparison of image compression with compaction

Table 1 compares the CASC method with conventional image storage and compression formats across three microscopy datasets with differing sparsity.

**Table 1.** Comparison of file size reduction achieved using conventional compression and content-aware compaction.

| **Specimen** | **TIFF** | **PNG** | **JPG** | **ZIP** | **JPG 2000** | **OME-ZARR** | **Data volume after CASC (OME TIFF)** | **Reduction in TIFF data volume after CASC** |
| --- | --- | --- | --- | --- | --- | --- | --- | --- |
| **DAPI-stained BPAE cell nuclei** | 37.4 MB | 1.14 MB | 965 kB | 1.17 MB | 1.52 MB | 1.8 MB | 731 kB | 51.2x |
| ***E. coli* JM105-miniTn7-*gfp*** | 8.20 MB | 2.84 MB | 62.2 kB | 2.56 MB | 2.3 MB | 3.65 MB | 21.0 kB | 390x |
| **Paxillin labelled in HeLa cells** | 238 MB | 26.1 MB | 7.81 MB | 26.2 MB | 18.5 MB | 35.4 MB | 12.6 MB | 18.9x |

Substantial reductions in file size were achieved following the application of CASC to all datasets studied. The greatest reduction was observed for the most sparse *E. coli* dataset, where the compacted output was reduced from 8.20 MB to 21.0 kB, corresponding to a 390-fold reduction in data volume. The BPAE nuclei dataset demonstrated a 51.2-fold reduction, while the denser paxillin-labelled HeLa cell dataset showed a more modest but still substantial 18.9-fold reduction.

As expected from Figure 7(B), the degree of reduction achieved by compaction varied according to the spatial sparsity of the input image. Datasets containing large background regions and isolated foreground structures exhibited the greatest reduction in file size, whereas denser datasets containing more continuous foreground occupancy showed lower but still significant reductions. This behaviour is consistent with the method explicitly removing empty spatial regions prior to reconstruction of the compacted image.

Compared with conventional lossless image formats including PNG, ZIP-compressed TIFF, JPEG2000, and OME-Zarr, the proposed method achieved greater reductions in stored data volume through spatial restructuring. In the *E. coli* dataset, the compacted image was smaller than all evaluated lossless formats, including PNG (2.84 MB), ZIP-compressed TIFF (2.56 MB), JPEG2000 (2.3 MB), and OME-Zarr (3.65 MB). Similar trends were observed for the BPAE nuclei dataset.

Lossy JPEG compression produced smaller file sizes than compaction for some datasets, particularly the *E. coli* images. However, unlike the proposed approach, JPEG compression alters pixel intensities and introduces irreversible compression artefacts. In contrast, the content-aware compaction method preserved object pixel intensities exactly while reducing spatial redundancy through rearrangement of foreground structures.

Because CASC intentionally sacrifices preservation of global spatial topology, the resulting file sizes are not directly equivalent to those produced by topology-preserving image compression methods. Instead, the results demonstrate the extent to which sparse microscopy datasets contain removable spatial redundancy that can be reduced through object-domain restructuring rather than conventional pixel-value compression alone.

### 3.8 Comparison with pixel-preserving object montages

To determine whether the observed reduction in spatial redundancy could be achieved using a simpler object montage representation, segmented foreground objects from each image dataset were assembled into a pixel-preserving montage. In contrast to conventional montages that resize objects to fixed display tiles, each object was retained at its original pixel dimensions and intensity values. Data are shown in Table 2. The montage representation preserved object dimensions and pixel intensities exactly but retained rectangular crop boundaries and associated background pixels.

**Table 2.**
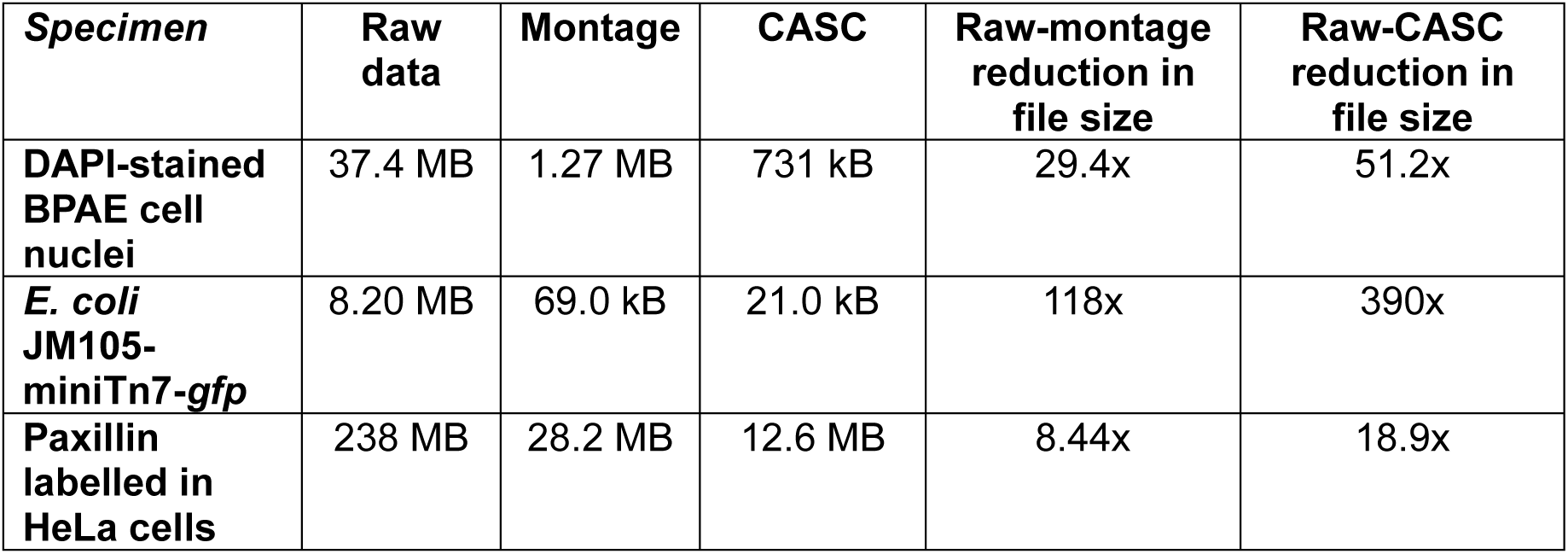
Comparison of file size reduction achieved using pixel-preserving montages and content-aware spatial compaction (CASC).

| ***Specimen*** | **Raw data** | **Montage** | **CASC** | **Raw-montage reduction in file size** | **Raw-CASC reduction in file size** |
| --- | --- | --- | --- | --- | --- |
| **DAPI-stained BPAE cell nuclei** | 37.4 MB | 1.27 MB | 731 kB | 29.4x | 51.2x |
| ***E. coli* JM105-miniTn7-*gfp*** | 8.20 MB | 69.0 kB | 21.0 kB | 118x | 390x |
| **Paxillin labelled in HeLa cells** | 238 MB | 28.2 MB | 12.6 MB | 8.44x | 18.9x |

Across all three datasets, CASC achieved greater reductions in file size than the corresponding pixel-preserving montage representation, reducing file size by 51.2x versus 29.4x for BPAE nuclei, 390x versus 118x for *E. coli*, and 18.9x versus 8.44x for paxillin-labelled HeLa cells. The montage required the retention of rectangular object crops together with inter-object spacing introduced by the shelf-based layout. In contrast, CASC reconstructs the compact image using object masks, allowing foreground structures to be packed more efficiently while preserving object pixel intensities.

These results demonstrate that preservation of object-level measurements is not unique to CASC. Rather, the principal advantage of CASC lies in its ability to combine quantitative object preservation with a more efficient reduction of spatial redundancy than can be achieved using a simple montage representation.

## 4. Conclusions

The CASC method presented here represents a departure from conventional approaches to image compression, which typically reduce redundancy in pixel values while preserving spatial layout. CASC explicitly restructures the spatial arrangement of foreground objects to remove empty image regions and reduce spatial redundancy. As such, CASC may be viewed not as a conventional image compression algorithm, but as a transformation from a spatially indexed image into a compact object-domain representation of the original data.

A key property of CASC is that object pixel intensities are copied directly from the original image during reconstruction. As a result, object-local intensity information and morphology are preserved without interpolation or lossy pixel transformation, making the method potentially suitable for downstream object-level quantitative analysis. At the same time, the reduction in spatial extent may enable faster visualisation and processing of sparse datasets.

A useful comparison can be made with pixel-preserving montage representations, in which segmented objects are extracted and rearranged into a compact layout while retaining their original dimensions and intensity values. Because such montages preserve object-local morphology and pixel intensities, preservation of object-level information is not unique to CASC. However, montage representations retain complete rectangular bounding boxes around each object, including local background pixels contained within the crop.

Comparison with pixel-preserving montages demonstrated that CASC consistently achieved greater reductions in data volume across all datasets examined. For the BPAE nuclei dataset, the montage reduced file size from 37.4 MB to 1.27 MB (29.4x), whereas CASC reduced file size to 731 kB (51.2x). For the *E. coli* dataset, the montage reduced file size from 8.20 MB to 69.0 kB (118x), whereas CASC reduced file size further to 21.0 kB (390x). Similarly, for the paxillin-labelled HeLa cell dataset, the montage reduced file size from 238 MB to 28.2 MB (8.44x), whereas CASC further reduced the file size to 12.6 MB (18.9x).

These results indicate that considerable reductions in data volume can be achieved simply by reorganising segmented objects into a montage representation. However, CASC consistently provided additional reductions beyond those achievable with montages. The magnitude of this advantage varied between datasets and was related to the proportion of background retained within the rectangular bounding boxes used by the montage representation. This improvement arises because CASC reconstructs the compact image using object masks rather than rectangular object crops, thereby eliminating background pixels retained within bounding boxes. Consequently, CASC provides a more efficient reduction of spatial redundancy while preserving object-local morphology and pixel intensities.

The quantitative spatial metrics demonstrated that the compaction process largely destroys both local neighbourhood relationships and global spatial ordering. This behaviour is an inherent consequence of the spatial repacking process rather than an implementation artefact. Accordingly, CASC is not suitable for applications in which absolute or relative spatial positioning is biologically meaningful, such as spatial transcriptomics (21), tissue architecture analysis (22), or studies of multicellular organisation. However, for many object-centric tasks, including object detection, classification, and morphology analysis, foreground objects may be treated largely independently (23), reducing the importance of preserving global spatial arrangement. In such cases, the reduction in spatial redundancy may outweigh the loss of spatial context.

The use of a heuristic shelf-based spatial packing strategy in CASC represents another limitation, as it does not guarantee optimal compaction. More advanced packing strategies could potentially yield tighter layouts, albeit at increased computational cost. Additionally, the practical performance of CASC is fundamentally dependent on the quality of foreground segmentation, since segmentation errors propagate directly into object extraction, packing, and reconstruction. The robustness experiments demonstrated that degradation in performance under severe noise or contrast reduction originated primarily from failures in segmentation rather than from the packing stage itself.

The present implementation of CASC used empirically selected parameters that provided stable performance across the microscopy datasets examined here, including an 8-pixel contextual padding margin and a minimum object size threshold of 20 pixels. These parameters are adjustable and may be modified depending on imaging modality, object scale, segmentation requirements, and the desired balance between contextual preservation and compaction efficiency. For example, increasing the padding margin preserves additional local background context but reduces packing efficiency, whereas smaller padding values increase compaction at the expense of contextual information. Future work could incorporate adaptive or data-driven parameter optimisation to improve robustness across diverse imaging conditions.

While demonstrated here for 2D microscopy images, the underlying concept is not inherently restricted to planar data. Extension of CASC to 3D and higher-dimensional imaging modalities represents a promising avenue for future research. Because sparsity often increases with dimensionality in biological imaging datasets, multidimensional compaction strategies may yield larger gains in storage efficiency, data transfer, and computational performance.

Beyond microscopy, the principles underlying this method may be applicable to other domains containing sparse image data, including satellite imaging, medical imaging, and point-cloud processing. In such applications, object-domain restructuring may provide a means of reducing spatial redundancy while preserving object-local information relevant to downstream analysis.

## Acknowledgements

I would like to thank Ainsley Beaton (University of Glasgow, UK) for the kind gift of the JM105-miniTn7-gfp strain. This work was funded in part by the Medical Research Council (MR/K015583/1), the Biotechnology and Biological Sciences Research Council (BB/Z51486X/1 and BB/X005178/1), and the Leverhulme Trust.

## Data availability

The data underlying this study are available from the University of Strathclyde Research Data Repository at DOI: 10.15129/c36e053d-887f-407b-ac74-462bdeb2f6e4. The dataset record is currently undergoing validation and will be publicly accessible upon publication.

